# Structure of a cytochrome-based bacterial nanowire

**DOI:** 10.1101/492645

**Authors:** David J. Filman, Stephen F. Marino, Joy E. Ward, Lu Yang, Zoltán Mester, Esther Bullitt, Derek R. Lovley, Mike Strauss

## Abstract

Electrically conductive pili from *Geobacter* species, termed bacterial “nanowires”, are intensely studied for their biological significance and potential in the development of new materials. We have characterized a unique nanowire from conductive *G. sulfurreducens* pili preparations by cryo-electron microscopy composed solely of the c-type cytochrome OmcS. We present here, at 3.4 Å resolution, a novel structure of a cytochrome-based filament and discuss its possible role in long-range biological electron transport.

**Summary sentence:** Cryo-electron microscopy reveals the remarkable assembly of a c-type cytochrome into filaments comprising a heme-based bacterial nanowire.

## Main text

Microbially produced protein nanowires are of profound interest due to their proposed role in global carbon and metal cycling and promise for myriad practical applications ^1–4^ Among the most thoroughly characterized are the electrically conductive pili (e-pili) of *Geobacter* species, which are comprised of pilin protein monomers ^5–7^ *Geobacter* e-pili enable long-range (μms) extracellular electron transfer to Fe(III) minerals and other microbial species, which is important in the biogeochemistry of diverse anaerobic soils and sediments ^3,8^. *Geobacter* species are among the most effective microbes in converting organic compounds into electricity because they can form thick electrically conductive e-pili biofilms ^9,10^. e-Pili purified from cells show potential as sustainably produced protein nanowires for electronics, as their properties can readily be tuned with genetic modifications to the pilin monomer; they can be incorporated into polymers to produce composite electronic materials; and they have dynamic sensing properties ^3,4,11^. Other bacteria produce e-pili from pilin monomers ^12,13^ and the archaeon *Methanospirillum hungatei* expresses an electrically conductive protein filament from archaellin monomers ^14^

Both e-pili and multi-heme outer-surface cytochromes have been implicated as being important for *Geobacter’s* long-range extracellular electron exchange with other cells and soil minerals^15,16^. Determining their respective contributions has been complicated by the previously reported association of OmcS, a cytochrome known to participate in some forms of extracellular electron exchange, with conductive filaments^17^. Immunogold labeling with anti-OmcS antibodies showed staining of thick filaments in pilus preps from *G. sulfurreducens*, leading Leang_ *et al*. to propose that OmcS may fulfil a functional role in directly modulating conductivity^18^. In pursuing investigation of these possibilities, we focused on the thick filaments in our e-pili preparations and have now solved the structure of these filaments by cryo-electron microscopy (cryo-EM). The 3D particle reconstructions of segments picked exclusively from thick filaments resulted in density that cannot be fit by any combination of PilA pilin monomers, but is perfectly fit by OmcS alone. We conclude that these thick filaments are not simply associated with OmcS but rather consist of OmcS monomers, thereby representing the first example of nanowires composed entirely of a bacterial cytochrome.

Preparations of conductive e-pili from *G. sulfurreducens* routinely contain multiple filaments with distinct morphological characteristics. In addition to the ca. 3 nm diameter filaments composed of the PilA pilin, thicker (ca. 4 nm) filaments are a prominent prep component (Fig. 1a). We pursued structural studies of these 4 nm, ostensibly OmcS-associated, filaments in an attempt to define their interaction with OmcS and determine its contribution to their conductive properties. Individual filament segments were extracted from cryo-EM images of conductive e-pili preps and, after analysis, the resulting 3D reconstruction produced a map with extensive contiguous polypeptide density that could not be satisfactorily fit with any combination of PilA monomers and also showed multiple regions of density along the filament providing excellent fits to heme molecules (Fig. 1b). In subsequent fitting attempts using the sequences of several related *Geobacter* cytochromes, only that of OmcS provided a near ideal fit to the density, incorporating the full OmcS sequence corresponding to the 47.5 Å filament repeat and accounting for all 6 hemes per OmcS monomer (Fig. 1c).

**Figure 1:**
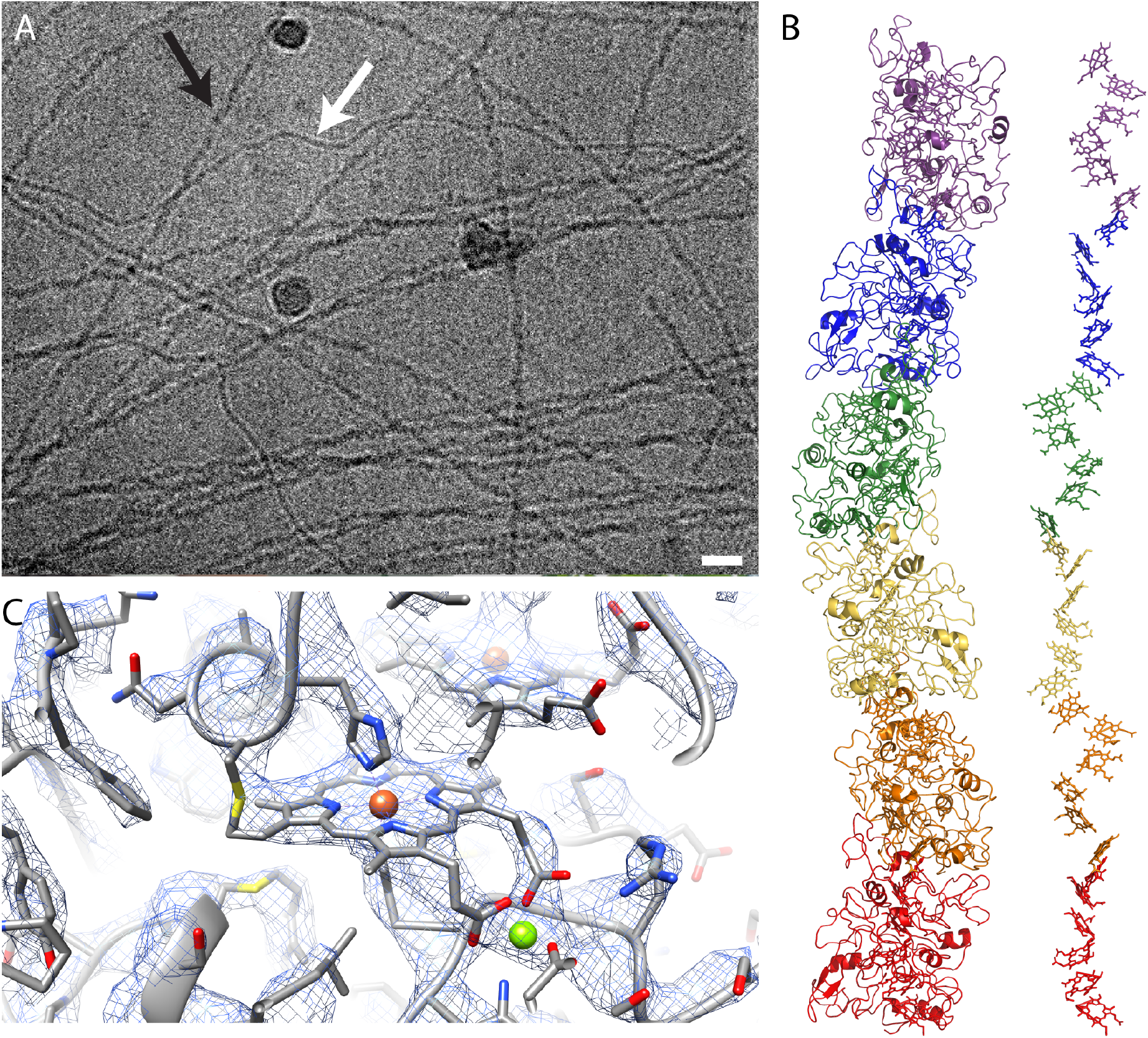
(A) Filaments isolated from Geobacter *sulfurreducens*, thicker OmcS filaments (black arrow), and thinner (3nm diameter) filaments (white) (Scale bar: 20nm). (B) Arrangement of monomers (left) and hemes (right) in filament. (C) Detail view of heme binding site.

The OmcS monomer from Geobacter sulfurreducens is a six-heme c-type cytochrome (UniProt Q74A86) and the OmcS fiber is a helically symmetric arrangement containing hundreds of monomers. Each monomer repeat is ~4.7nm along the fiber axis, rotated 83 degrees and ~4nm wide, except at the inter-monomer interfaces, where the fiber narrows significantly (Fig. 2a,b). At the core of the fiber is a continuous, unbranched stack of hemes (Fig. 1c, 2c).

**Figure 2:**
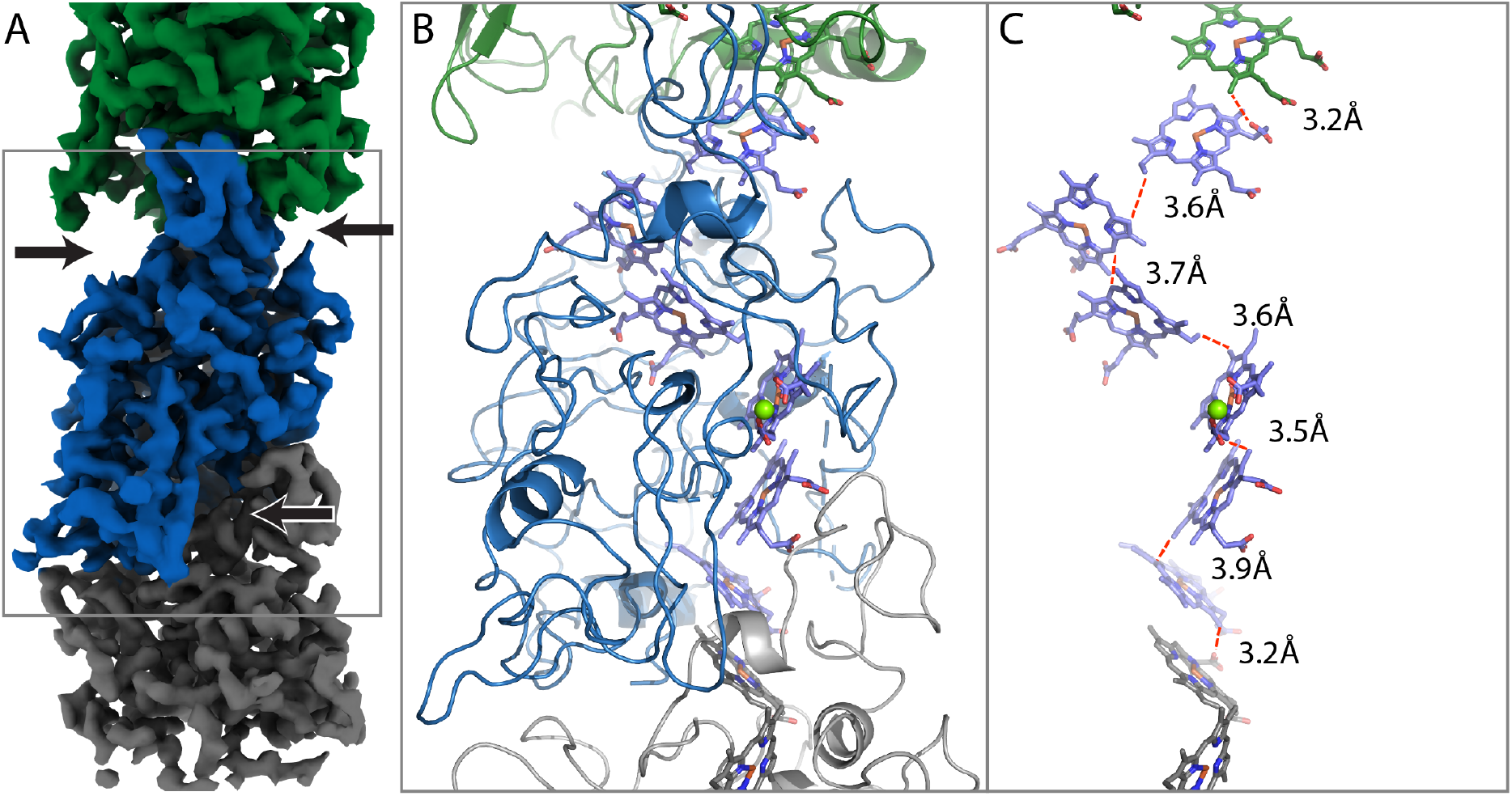
Filament monomer as surface rendering (A) and model view (B), with arrows pointing to cut-outs. (C) Interatomic contact distances between adjacent porphyrins are 3.9A or less.

Each heme has two nearest neighbours, one above and one below. Successive hemes, approaching edge-to-edge, alternate between lying parallel and perpendicular to the previous heme (Fig. 2c). The interatomic contact distances are all ≤ 3.9Å (see Supp Table 1). This proximity suggests a mechanism for long-range electrical conduction, but leaves open the question of how charges enter and exit the nanowire.

The OmcS sequence includes six Cys-x-x-Cys-His motifs, with both Cys sulfur atoms covalently attached to heme substituents, and His as one of the axial ligands (Fig 1c). The stacking order of hemes along the fiber axis coincides with the order of the Cys-x-x-Cys-His motifs in the sequence. This exact correspondence has been seen in some (e.g., PDB 3ov0^19^), but not all (e.g PDB 1z1n), multi-heme cytochromes. All 12 heme-axial ligands are histidine side chains, with 11 coming from the same OmcS subunit, and one (attached to heme959) from a neighbouring molecule.

The OmcS protein is well-ordered along its nearly continuous main chain, with distinct bends at most of its alpha carbon positions (Fig 1c) and side chain densities consistent with the amino acid sequence, except for 15 poorly ordered residues that were truncated in the model. The protein wraps around the central heme stack, nearly completely shielding the hemes from solvent. Notable exceptions, which are solvent-exposed edge-on, include heme929 (from one OmcS subunit) and heme969 (from the adjacent subunit), which are both located near the narrow, inter-subunit interfaces (on opposite sides of the fiber). These two sites could have a function in conduiting electrons into or out of the filament.

The heme stack is integral to both the monomer and fiber structures, and dictates the folding pattern of the surrounding 407 residue protein chain, resulting in remarkably little regular secondary structure, with fewer than 20 residues as alpha helices using KSDSSP^20^, and the overall fold is unique among solved protein structures (by VAST search, https://www.ncbi.nlm.nih.gov/Structure/VAST/vastsearch.html).

Each OmcS monomer also includes one octahedrally coordinated metal, tentatively identified as Mg^++^ based on its density level and experiment (see Supp. Table 2), whose biological significance is unclear. Two of its ligand positions are occupied by acidic substituents of heme949 (see Fig 1c).

The unexpected filamentous assembly of a *c*-type cytochrome and our structure suggests new possibilities for long-range biological electron transfer. Heme-to-heme electron transport occurs within individual multi-heme *c*-type cytochromes and electron transfer between two multi-heme cytochromes held within a barrel protein is possible^21^, though nanowires comprised solely of multiple cytochromes have not been observed. The cytochrome-rich membrane extensions of *Shewanella oneidensis*, when dried, are electrically conductive ^22,23^. However, under physiologically relevant hydrated conditions, gaps between the membrane associated cytochromes prevent long-range cytochrome-to-cytochrome electron hopping/tunneling ^1,24^.

Several lines of evidence attest to the importance of OmcS on the extracellular surface of *Geobacter* cells. It is present on the outer membranes of both wild-type cells^18^ and cells unable to produce e-pili^25^. OmcS deletion strains are deficient in Fe(III) oxide reduction but expression *in trans* rescues the phenotype^26^, and strains overexpressing OmcS show enhanced electron transfer between *Geobacter* species^27^. This suggests that OmcS filaments may increase the coverage of the cell surface with redox-active moieties beyond that possible with other surface cytochromes that are connected to the periplasm through outer-membrane conduits ^16^. OmcS filaments may also provide an electron transfer path between surface cytochromes electrically connected to the periplasm and e-pili for long-range extracellular electron transfer. Additional studies are warranted to determine if nanowires comprised of hemes are present in other microbes and to explore the possibilities of employing heme-based nanowires as a sustainably produced material for electronic devices.

## Materials and Methods

### Preparation of cryo-EM samples, data collection and processing

Nanowires were collected and purified as previously described in Tan *et al* ^28^ and then dialyzed against deionized water to remove buffer constituents. Samples for cryoEM were added to glow discharged R2/2 Quantifoil grids and plunge-frozen on a Vitrobot Mk IV (ThermoFisher). Imaging was carried out on a Titan Krios (ThermoFisher) equipped with a Bioquantum K2 GIF (Gatan Inc) using a magnification corresponding to a 1.06Å calibrated pixel size. The 45-frame acquisitions were acquired with SerialEM using counting mode and accumulating a total dose of 50 e^−^/Å^2^ over 15 seconds. Frame alignment was done in MotionCor2, and all subsequent processing steps were carried out in Relion-3^29^. Filaments were manually selected and overlapping segments were extracted, and refined to produce a 3.9Å map. 2D and 3D classification was carried out, but proved unnecessary, due to the homogeneity of the filaments selected. After particle polishing, the final reconstruction was consistent with a 3.4Å map (as determined by FSC 0.143; Supp. Fig. 1).

### Model building and refinement

Early inspection of the reconstruction clearly showed a line of metal sites running along the fiber axis, and a continuous stack of six porphyrins per helical subunit. Detailed atomic models for the protein were added to the reconstruction, repeatedly rebuilt using COOT^30^ and SPDBV^31^, and refined using Refmac5^32^ and Phenix.autobuild^33^. Initial efforts to find one or more copies of the 61-amino acid protein PilA in the map never yielded a convincing fit and several density patterns were identified that were incompatible with the PilA sequence. To discover what else might be present, atomic models were prepared with arbitrary sequences closely resembling the density. Once sequence database searches, using psi-Blast^34^, found similarities to a number of related six-heme C-type cytochromes, we focused, among these, upon OmcS, which was known to be abundant in the fiber sample. An atomic model with the OmcS sequence was fitted to the map, and produced a convincing visual fit.

In Refmac5, a stereochemically restrained atomic model, with RMS bond length deviations of 0.012.Å, and overall temperature factors, yielded an R-factor of 32.2% (versus pseudo-crystallographic data in space group P4(3)), and an average Fourier shell correlation of 0.68. Submitted coordinates were then rerefined versus the authentic map.

## Supporting information

## Acknowledgements

This research was supported by Office of Naval Research grant N00014-16-1-2526. Accession number: EMDB XXXX, PDB YYYY

## References

1. Ing, N.L. et al. J. Phys. Chem. B, DOI: 10.1021/acs.jpcb.8b07431 (2018).

2. Creasey, R.C.G. et al. Acta Biomater. 69, 1–30 (2018).

3. Lovley, D.R. Curr. Opin. Electrochem. 4, 190–198 (2017).

4. Lovley, D.R. mBio 8, e00695–17 (2017).

5. Reguera, G. et al. Nature 435, 1098–1101 (2005).

6. Malvankar, N.S. et al. mBio 6, e00084–15 (2015).

7. Holmes, D.E. et al. Microbial Genomics 2, doi: 10.1099/mgen.0.000072 (2016).

8. Lovley, D.R. Ann. Rev. Microbiol. 71, 643–664 (2017).

9. Malvankar, N.S. et al. Nat. Nanotech. 6, 573–579 (2011).

10. Vargas, M. et al. mBio 4, e00105–13. (2013).

11. Sun, Y.-L. et al. Small 14, 1802624 (2018).

12. Walker, D.J.F. et al. ISME J. 12, 48–58 (2018).

13. Walker, D.J.F. et al. bioRxiv (2018).

14. Walker, D.J.F. et al. bioRxiv, doi: https://doi.org/10.1101/458356 (2018).

15. Lovley, D.R. Annu. Rev. Microbiol. 71, 643–664 (2017).

16. Shi, L. et al. Nat. Rev. Microbiol. 14, 651–662 (2016).

17. Tan, Y. et al. Front. Microbiol. 7, 980 (2016).

18. Leang, C. et al. Appl. Environ. Microbiol. 76, 4080–4084 (2010).

19. Pokkuluri, P.R. et al. J. Struct. Biol. 174, 223–33 (2011).

20. Kabsch, W. & Sander, C. Biopolymers 22, 2577–637 (1983).

21. Blumberger, J. Curr. Opin. Chem. Biol. 47, 24–31 (2018).

22. Pirbadian, S. et al. Proc. Natl. Acad. Sci. U. S. A. 111, 12883–12888 (2014).

23. El-Naggar, M.Y. et al. Proc. Natl. Acad. Sci. USA 107, 18127–18131 (2010).

24. Subramanian, P. et al. Proc. Nat. Acad. Sci. 115, E3246–E3255 (2018).

25. Flanagan, K.A. et al. bioRxiv, 114900 doi: https://doi.org/10.1101/114900 (2017).

26. Mehta, T. et al. Appl. Environ. Microbiol. 71, 8634–8641 (2005).

27. Summers, Z.M. et al. Science 330, 1413–1415 (2010).

28. Tan, Y. et al. mBio 8, e02203–16 (2017).

29. Zivanov, J. et al. eLife 7(2018).

30. Emsley, P. & Cowtan, K. Acta Crystallogr. D Biol. Crystallogr. 60, 2126–32 (2004).

31. Guex, N. & Peitsch, M.C. Electrophoresis 18, 2714–23 (1997).

32. Winn, M.D. et al. Methods Enzymol. 374, 300–21 (2003).

33. Zwart, P.H. et al. Methods Mol. Biol. 426, 419–35 (2008).

34. Altschul, S.F. et al. Nucleic Acids Res. 25, 3389–402 (1997).

