## Supplementary material for "Structure of a cytochrome-based bacterial nanowire"

### Supplemental Table 1

In Supplemental Table: Closest interatomic contacts between neighbouring hemes.

| Heme | Atom | Heme | Atom | Distance (Å) |
|---|---|---|---|---|
| 919 | CBC | 929 | CBB | 3.6 |
| 929 | CHB | 939 | CHB | 3.7 |
| 939 | CBC | 949 | CMC | 3.6 |
| 949 | C3A | 959 | CMA | 3.5 |
| 959 | CBC | 969 | CHC | 3.9 |
| 969 | CMA | 919 | O1A | 3.2 |

### Supplemental Table 2: Inductively Coupled Plasma Mass Spectrometry

| | Concentration, mg/kg | | | | | | | | | | | | | |
|---|---|---|---|---|---|---|---|---|---|---|---|---|---|---|
| **Sample Name** | **Fe** | **SD (Fe)** | **Mg** | **SD (Mg)** | **K** | **SD (K)** | **Ca** | **SD (Ca)** | **Mn** | **SD (Mn)** | **Zn** | **SD (Zn)** | **Sr** | **SD(Sr)** |
| Sample W | <0.005 | | <0.01 | | <0.02 | | <0.01 | | <0.002 | | <0.002 | | | <0.002 |
| Sample B | 0.049 | 0.005 | <0.01 | | 0.12 | 0.01 | <0.01 | | 0.0029 | 0.0003 | <0.002 | | | <0.002 |
| Sample B+W | 0.104 | 0.003 | <0.01 | | <0.02 | | <0.01 | | 0.0028 | 0.0003 | <0.002 | | | <0.002 |
| Sample G Sulf Pili | 1.47 | 0.02 | 0.61 | 0.02 | 0.41 | 0.02 | 0.24 | 0.01 | 0.0125 | 0.0006 | 0.01 | 0.001 | 0.0046 | 0.0005 |

sample w = milli-Q water directly from our system
sample b = 150mM Ethanolamine, pH 10.5 made using Ethanolamine-HCl plus NaOH.
sample w + b = 150mM Ethanolamine, pH 10.5 dialyzed against milli-Q water

**Supplemental Figure 1**

Fourier Shell Correlation for cryoEM reconstruction of OmcS filament.

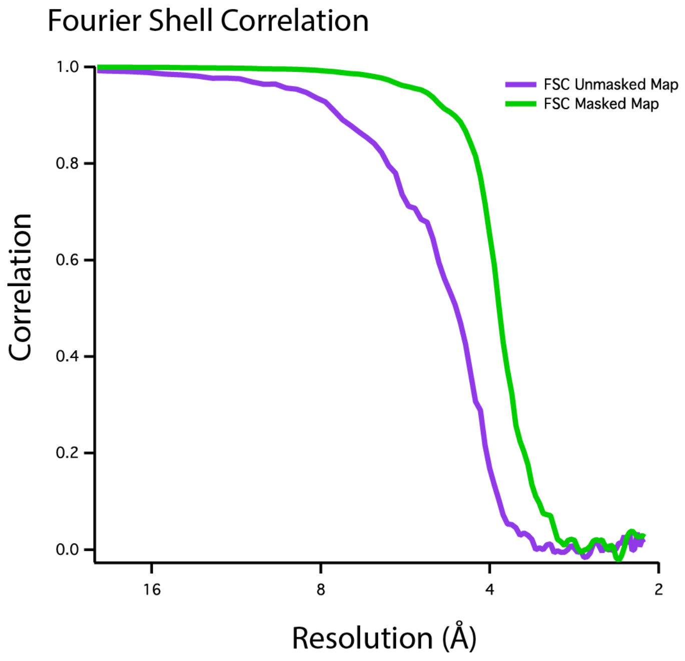
